# Floating Sphere Assay: A rapid qualitative method for microvolume analysis of gelation

**DOI:** 10.1101/2022.03.21.485096

**Authors:** Uma Jingxin Tay, Megan Goh, Jeralyn Ching Wen Hui, Prakash Arumugam

**Affiliations:** Singapore Institute of Food and Biotechnology Innovation, 61 Biopolis Drive, Singapore 138673; Nanyang Technological University, School of Biological Sciences, Singapore

**Keywords:** Gelation screen, minimum gelling concentration, Inversion Assay, Floating Sphere Assay

## Abstract

1

A huge, unprecedented demand for gelatin coupled with its implications on global sustainability has resulted in the need to discover novel proteins with gelling attributes for applications in the food industry. Currently used gelation assays require large sample volumes and thus the screening for novel gelling proteins is a formidable technical challenge. In this paper, we report the ‘Floating Sphere Assay’ which is a simple, economical, and miniaturized assay to detect minimum gelling concentration with volumes as low as 50 μl. Results from the Floating Sphere Assay are consistent with currently used methods for gelation tests and accurately estimate the Minimum Gelling Concentrations (MGCs) of gelatin, κ-carrageenan and gellan gum. The assay was also able to differentiate the strengths of strong and weak gellan gum gels prepared at pH 3.5 and pH 7.0 respectively. The Floating Sphere Assay can be utilized in high-throughput screens for gelling proteins and can accelerate the discovery of gelatin substitutes.

**Highlights:**

- We report the Floating Sphere Assay that can be used to assesses minimum gelling concentration of solutions with volumes as low as 50 μl.
- Observing whether a glass sphere placed on the surface of a test solution floats or sinks is diagnostic of gel formation
- Floating Sphere Assay can distinguish a strong gel from a weak gel
- Floating Sphere Assay is a rapid and cost-effective approach to screen for novel plant-based gelatin alternatives.

## 4 Introduction

Gelling agents are used in a wide variety of convenient foods including pastas, salad dressings, yoghurt, ice-creams, jams, low calorie meals as well as restructured foods. A large proportion of gelling agents are polysaccharides like pectin, Sodium alginate, carrageenan and gellan gum. Gelatin is the only member of the protein family that is used in food gels. Polysaccharides are soluble dietary fibers and thus would be fermented by gut bacteria. Excessive fermentation leads to undesirable side effects such as flatulence, bloating, stomach discomfort, diarrhea and constipation (Makki, Deehan, Walter, & Bäckhed, 2018). Consequently, gelatin is a preferable alternative for individuals prone to irritable bowel syndrome and colitis. Apart from being a widely used food ingredient, gelatin is increasingly adopted to constitute cell culture scaffold in engineering cell-cultured meat by virtue of its biocompatibility, biodegradability and low immunogenicity (MacQueen, et al., 2019). Consequently, the high reliance on gelatin which already has a market volume of 620.6 kilotons in 2019, is expected to continue growing at a volume-based Compound Annual Growth Rate of 5.9% between 2020 to 2027 (Grand View Research., 2020).

Present reliance on gelatin is however unsustainable. Gelatin is mainly sourced from bones, skins, and tendons of bovines and pigs and to a lower extent from fish and poultry (Grand View Research., 2020). However, animal farming is inherently inefficient, straining the environment, public health and food security. Moreover, gelatin is not halal. Thus, the need for identifying plant-based protein alternatives for gelatin cannot be exaggerated.

It is possible to envisage two kinds of screening strategies for gelling proteins. In the first ‘activity-guided approach’, protein extracts from food-grade systems are fractionated and the gelling ability of individual protein fraction is assessed. In the second ‘candidate protein approach’, specific proteins from food-grade systems are expressed in hosts such as *E. coli* and purified, and the gelling ability of purified protein solutions is tested. Gelatin forms gels in the concentration range of about 20-100 mg/ml (Djabourov, Lechaire, & Gaill, 1993; Pandey, Tiwari, Pandey, Rao, & Das, 2019). Assuming a similar concentration range for gelatin substitutes, large amounts of proteins would be required for performing such screens. Moreover, it will be necessary to sample a range of buffer and pH conditions for gelation screens. In the ‘activity-guided approach’, the gelling abilities of only the highly expressed proteins would be feasible to test and even this will require concentrating the protein fractions to small volumes. In the ‘candidate protein approach’, efficient expression and purification of the protein is essential. It is therefore highly desirable to develop miniaturized assays that require minimal solution volumes for assessing gelation.

The most widely used qualitative assay for gelation is the so-called ‘Inversion Assay’. In this assay, the tube containing a given sample is inverted to test if the solution flows or not (Figure 1a). If the solution does not flow, it is considered to have formed a gel. If the solution flows, it would indicate absence of gelation.Minimum Gelling Concentration (MGC) corresponds to the lowest concentration of the gelling agent at which the solution stays firm when tubes are inverted (Aydemir & Yemenicioğlu, 2013; Yang, et al., 2018). However, this assay is not suitable for gelation screens with small solution volumes. For example, 500 μL solutions (non-gels) were observed to resist flow in 1.5 ml tubes (Hughes, et al., 2018). As solutions resist flow presumably due to surface tension and adhesive forces between tube and the molecules, gelation tubes require a wide mouth, necessitating large sample volumes for the assay.

**Figure 1.**
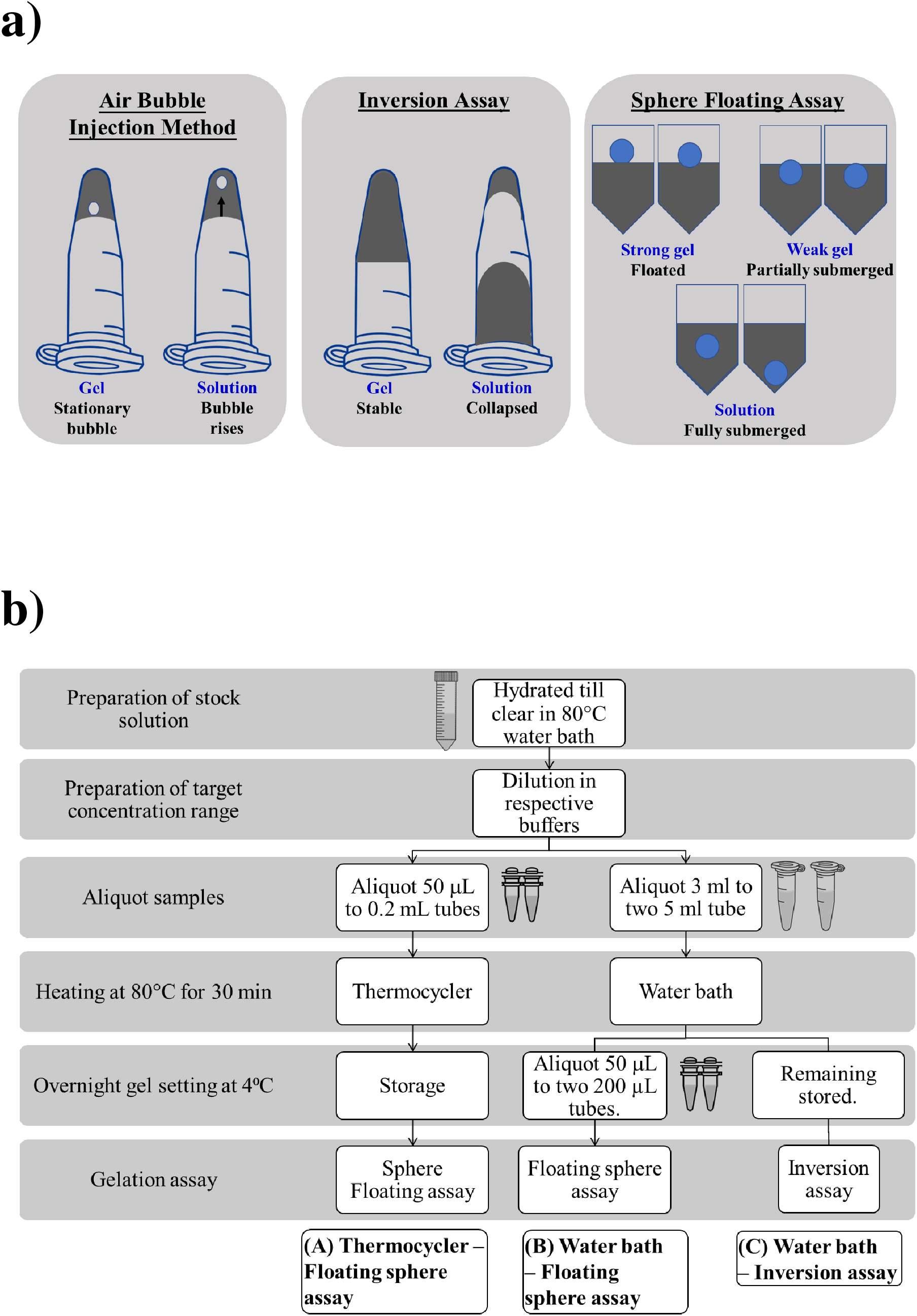
Floating Sphere Assay for assessing gelation. (a) Comparison of the proposed Floating Sphere Assay against conventional approaches; (b) Outline of the experimental strategy for Floating Sphere Assay.

As an alternative to the Inversion Assay, Hughes, et al. (2018) used an air bubble injection method – a bubble is introduced to samples (~500 μl) in inverted 1.5 ml microfuge tubes and gelation is detected by tracking the bubble. If the bubble rises to the top of the tube, it is a solution and if the bubble remains at the liquid surface, it is a hydrogel (Figure 1a). However, injecting a bubble without fracturing the gel will be a challenge for brittle hydrogels. Fragmentation of the gel would be particularly pronounced with small sample volumes. Moreover, tracking the movement of the air bubble might not be straightforward.

In this paper, we report a simple miniaturized gelation assay referred to as the ‘Floating Sphere Assay’ that requires sample volumes as low as 50 μl (Figure 1a). This method is adapted from the viscometrical falling sphere assay that determines viscosity of solutions based on terminal velocity measurements of a ball dropped into the solution via Stokes equation (Ali, Al-Zuky, Al-Saleh, & Mohamad, 2019; J. X. Tang, 2016). On dropping the ball onto the solution, we assessed whether it floats on top (i.e. less than 25% of the bead has penetrated into the sample); partially submerges (more than 25% of the bead has penetrated) or fully submerges (the bead has sunk below the surface of the sample).

The Floating Sphere Assay can accurately determine the MGC of widely used gelling agents namely gelatin, potassium-supplemented κ-carrageenan and calcium-supplemented low acyl gellan. Gelatin is dominated by repeating motifs or glycine-(hydroxyproline or proline)-proline.

On cooling, gelatin partially returns from disordered coils to polyproline II helix, resembling that of collagen. Around three regions of the helix generally interact. These regions transit from intra to inter-helix at higher concentrations, forming junction zones that induce gelation (Pang, Deeth, Sopade, Sharma, & Bansal, 2014). In contrast, gellan and κ carrageenan are linear anionic polysaccharides. Gellan comprises of tetrasaccharide repeats of [→3)-β-D- glucose- (1→4)-β-D-glucuronic acid −(1→4)-β-D-glucose-(1 → 4)-α-L-rhamnose-(1→)]. On cooling, gellan transits from disordered coils to double helices. Removal of acetyl substituents in the low acyl variant greatly reduces entropic and steric hindrance of the low acyl gellan gum helices to aggregation which is required for gelation. Aggregation of the double helices can be promoted by reducing electrostatic repulsion between glucuronate carboxylate groups through introducing cations (e.g. calcium) or lowering the pH. When calcium is incorporated, coordination binding between calcium and gellan gum further facilitates aggregation of the double helices (Morris, Nishinari, & Rinaudo, 2012). κ-carrageenan comprises of disaccharide repeats of [→3)-β -D-galactose 4-sulfate-(1→4)-3,6-anhydro-α-D-galactose-(1→)]. Like gellan gum, κ-carrageenan undergoes coil-helix transition in the presence of potassium which coordinate the sulphate groups of κ-carrageenan, and aggregate into gel networks (Chen, Liao, & Dunstan, 2002).

Overall, the results of gelatin, potassium κ-carrageenan and calcium-low acyl gellan from Floating Sphere Assay show good concordance with results from the widely used Inversion Assay. Moreover, the Floating Sphere Assay was able to distinguish the strength of strong and weak gellan gum gels prepared at pH 3.5 and 7.0 respectively.

## 5 Material and methods

### Preparation of gels

Stock solutions of the gelling agents were prepared by hydrating the respective powders in their corresponding gelation buffers prepared using Milli-Q Water. Calcium κ-carrageenan was prepared by hydrating κ-carrageenan (22048, Sigma-Aldrich, Germany) in 200 mM KCl solution to form a 2% solution (Chen, et al., 2002; Hermansson, Eriksson, & Jordansson, 1991). Calcium-gellan gum was prepared by hydrating low acyl gellan gum (Kelcogel, CP Kelco, USA) in 5 mM CaCl_2_ to form a 2% solution (Morris, et al., 2012). Whereas gellan gum of pH 3.5 and 7.0 were prepared by hydrating low acyl gellan gum in respective buffers of pH 3.5 ± 0.1 and 7.0 ± 0.1 to form a 2% solution. Buffers were made by adjusting Milli-Q water using 5 M NaOH and 80% L-(+)-Lactic acid (27715, Sigma-Aldrich, Germany). Gelatin was prepared by hydrating porcine gelatin (G1890, Sigma-Aldrich, Germany) to form a 5% solution. Stock solutions were prepared by heating in an 80°C water bath under shaking at 90 rpm (SW22, JULABO GmbH, Germany) (Chen, et al., 2002; Morris, et al., 2012). The corresponding gelation buffer was similarly heated in the 80°C water bath and used for diluting the stock solution. For gelatin, a concentration range of 0.625% – 5.0% (0.625%, 1.25%, 2.50% and 5.0%) was used. For all other gelling agents, a concentration range of 0.0625 – 2.0% (0.0625%, 0.125%, 0.25%, 0.50%, 1.0%, 1.5% and 2.0%) was used.

For each specified concentration of the gelling agent, 3 ml was aliquoted to each of the duplicate 5.0 mL tubes (Eppendorf Tubes^®^, Germany) while 50 μL was aliquoted to each of the duplicate 200 μL Polymerase Chain Reaction (PCR) tubes (Axygen^®^, Corning Incorporated, USA). Samples in the 5 mL tubes and PCR tubes were respectively heated at 80°C for 30 minutes using the water bath and thermocycler (SimpliAmp™ Thermal Cycler, Applied Biosystems™, ThermoFisher Scientific). After the heating step, 50 μL of each solution was immediately aliquoted from the 5.0 mL tube to each of the duplicate 200 μL tubes. Phosphate buffered saline (PBS) was also included in the PCR tubes as a negative control. All tubes were stored at 4°C overnight. On the following day, tubes were left to equilibrate to room temperature of around 24 °C for an hour prior to performing the gelation assay. Biological triplicates were performed in comparing the results of Inversion Assay with the Floating Sphere Assay. The procedure is summarised in Figure 1b.

### Floating sphere Assay

Borosilicate glass beads of 3.0 mm (Z143928, Merck KGaA, Germany) were coated with blue ink from Sharpie Permanent Marker Fine Point (Newell Office Brands, USA). The permanent marker was pierced to obtain the ink cartridge. Ink from the cartridge was gently pressed out into a petri-dish. Glass beads were added to the petri dish which was then sealed and shaken vigorously to coat the glass beads evenly with the ink. Density of glass beads was averaged from the mass and radius (Mitutoyo Digimatic Micrometer, Japan) of 10 glass beads assuming spherical dimensions. Using a tweezer, beads were deposited on to the top of the gelling solution. After 5 minutes, the position of the beads was scored as either floating or partially submerged or fully submerged.

## 6 Results and Discussion

### A simple miniaturized assay for gelation

Our motivation to develop the Floating Sphere Assay was driven by our observation that inverting 200 μL tubes containing 50 μL of PBS did not result in the solution flowing to the bottom (Supplementary Figure 1). For microvolumes such as 50-500 μl, the Inversion Assay is therefore not suitable to detect MGC. Consequently, we developed a gelation assay based on the falling sphere assay which is commonly used to determine viscosities of solutions. By measuring the terminal velocity of a sphere dropped into the solution, the viscosity of the solution can be determined using the Stokes equation. Lower the terminal velocity of the sphere, higher the viscosity of the solution. For our assay, we chose glass beads that have a higher density (2.64 g/cm^3^) than commonly used gelling aqueous solutions (~1 g/cm^3^) to ensure that weight of the sphere overcomes the buoyant forces. We reasoned that if a gel is formed, it might be able to counter the weight of the sphere and thereby prevent it from entering the ‘solution phase’.

To test this above idea, we chose three gelling agents namely gelatin, gellan gum and κ-carrageenan and compared the results obtained with the traditional Inversion Assay. We prepared solutions containing each of the gelling agent at different concentrations in 5 mL tubes and heated the solutions at 80 °C in a water bath for 30 min (Figure 1b). To evaluate the possibility of using smaller volumes for gelation, on completion of the heating step, we transferred a small aliquot of the solution from the 5 ml tube into a 0.2 mL tube and then incubated both 5 mL and 200 μL tubes at 4 °C overnight to promote gelation. For screens of gelling proteins, it will not be possible to use the water bath for heating since this requires tubes larger than 200 μL but small tube volumes are required to minimise sample volumes. Therefore, to evaluate the potential of heating 200 μL tubes directly, we assessed whether thermocycler is as effective as water bath in heating the gelling agent solutions. In parallel, we prepared 50 μL of the gelling agent solutions in 200 μL tubes and performed the heating step in thermocycler and subsequently stored at 4 °C overnight for the gels to set (Figure 1b). After overnight incubation, we equilibrated tubes to room-temperature. Inversion Assay was performed using samples in 5 mL tubes and the Floating Sphere Assay with samples in 200 μ L tubes.

Overall, we saw a very good correlation between results from the Inversion and the Floating Sphere Assays for all the three gelling agents. When the bead floated or was partially submerged in the solution in Floating Sphere Assay, gelation was always observed in the Inversion Assay. Solutions in which the bead fully submerged, also did not form gels in the Inversion Assay. The only exception occurred for pH 7 gellan gum where at 0.5%, the bead remained partially submerged in the Floating Sphere Assay but the sample had collapsed in the Inversion Assay. Results from our experiments are presented in Tables 2 – 4.

**Table 1.**
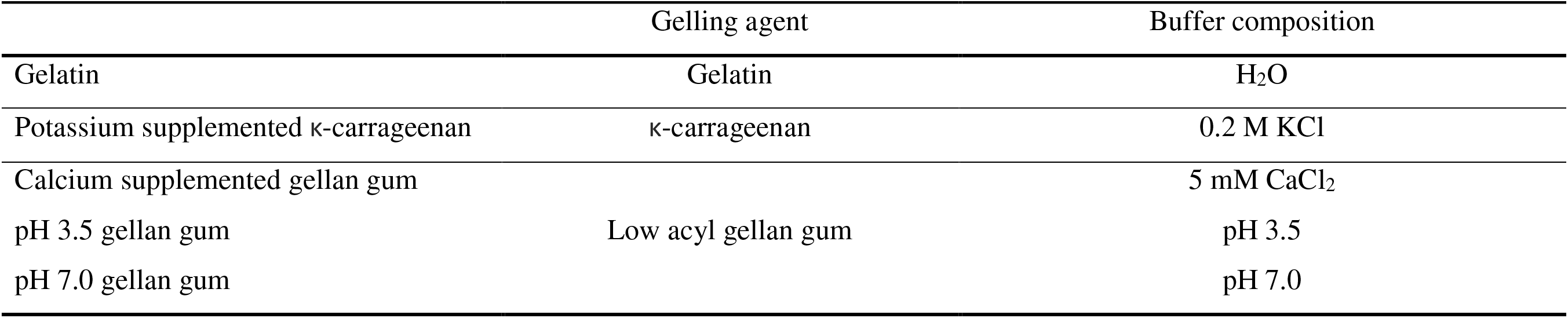
Summary of gelling agents evaluated.

**Table 2.** Gelation results of gelatin.

|  | Concentration (%) |  |  |  |
| --- | --- | --- | --- | --- |
|  | 5.0 | 2.5 | 1.25 | 0.625 |
| (A) Thermocycler – Falling sphere assay | Float | Float and PS | PS | FS |
| (B) Water bath – Falling sphere assay | Float | Float | PS | FS |
| (C) Water bath – Inversion assay | Stable | Stable | Stable | Collapsed |

**Table 3.** Gelation results of potassium-supplemented κ-carrageenan.

|  | Concentration (%) |  |  |  |  |  |  |
| --- | --- | --- | --- | --- | --- | --- | --- |
|  | 2.0 | 1.5 | 1.0 | 0.5 | 0.25 | 0.125 | 0.0625 |
| (A) Thermocycler – Falling sphere assay | Float | Float | Float and PS | Float and PS | PS | PS | FS |
| (B) Water bath – Falling sphere assay | Float | Float | Float and PS | Float and PS | PS | PS | FS |
| (C) Water bath – Inversion assay | Stable | Stable | Stable | Stable | Stable | Collapsed | Collapsed |

**Table 4.** Gelation results of calcium-supplemented low acyl gellan gum.

|  |  | Concentration (%) |  |  |  |  |  |  |
| --- | --- | --- | --- | --- | --- | --- | --- | --- |
|  |  | 2.0 | 1.5 | 1.0 | 0.5 | 0.25 | 0.125 | 0.0625 |
| (A) Thermocycler – Falling sphere assay |  | Float | Float | Float | Float and PS | PS | PS | FS |
| (B) Water bath – Falling sphere assay |  | Float | Float | Float and PS | Float and PS | PS | PS | FS |
| (C) Water bath – Inversion assay |  | Stable | Stable | Stable | Stable | Stable | Stable | Collapsed |

Gelatin formed a gel at a minimum gelling concentration of 1.25% in both the Floating Sphere and Inversion Assays. Representative images of gelation assays of gelatin (Table 2a) are presented in Figure 2 (Data from two additional biological replicates are presented in Supplementary Figure S2). For 1.25 – 5.0%, the bead either floated or was partially submerged in the Floating Sphere Assay and did not flow in the Inversion Assay. Whereas, in the 0.625% solution of gelatin, the bead was fully submerged in the Floating Sphere Assay and collapsed in the Inversion Assay.

**Figure 2.**
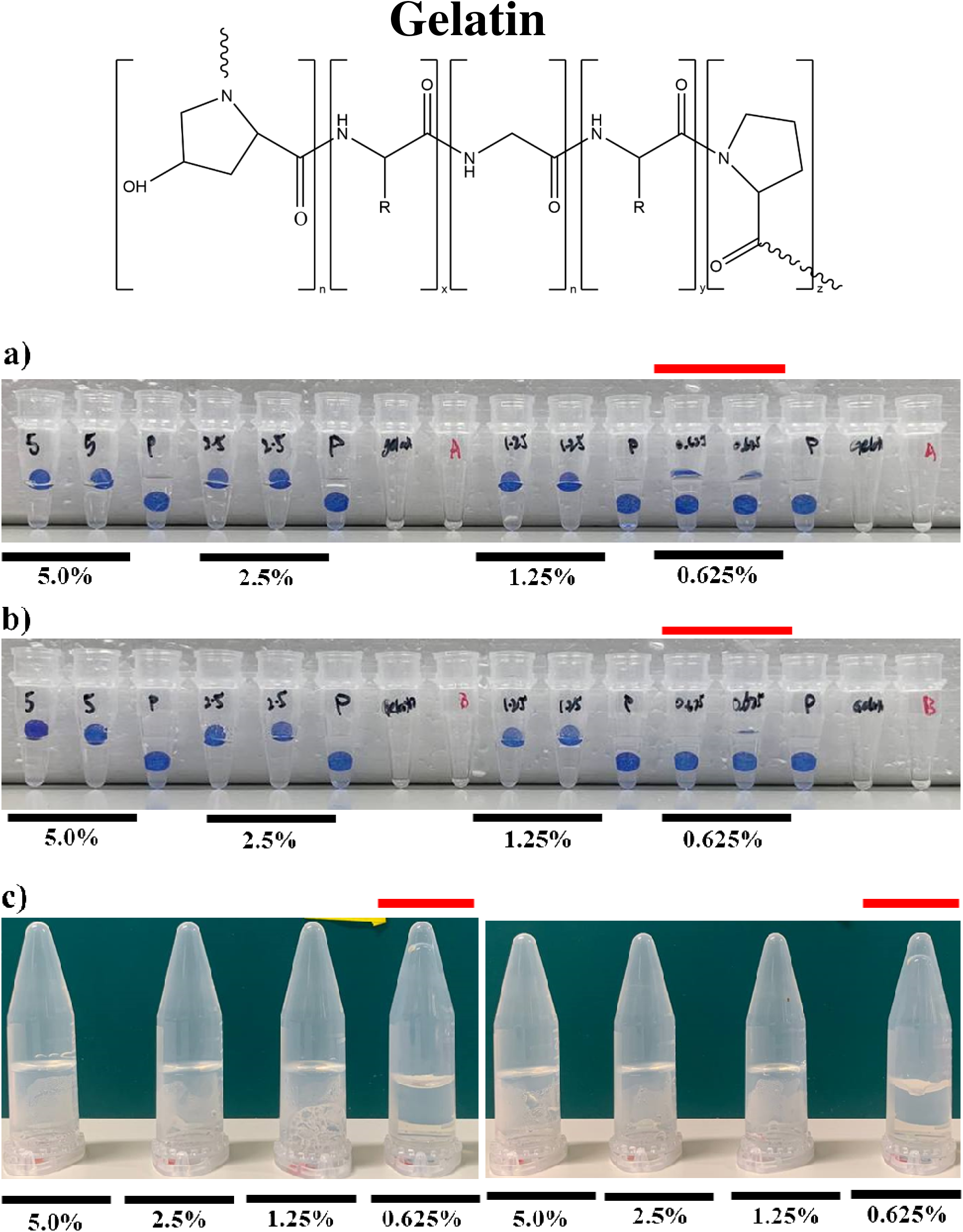
Determining the Minimum Gelling Concentration of gelatin via the Floating Sphere and Inversion Assays. Representation of gelation results of gelatin in (a) Thermocycler – Floating Sphere Assay; (b) Water bath – Floating Sphere Assay and (c) Water bath – Inversion Assay. Red line indicates the concentrations below the critical gelling concentration. A minimum of three biological replicates was performed. Results from two additional replicates are presented in Supplementary Figure S2.

Similar trends were obtained for potassium κ-carrageenan (Table 3) and calcium-supplemented low acyl gellan gum (Table 4) and the data are presented in Figure 3 and 4 respectively (Data from two additional biological replicates are presented in Supplementary Figures S3 and S4). For 0.125 – 2.0% solutions, the bead either floated or was partially submerged in the Floating Sphere Assay and did not flow in the Inversion Assay. Whereas, in the 0.0625% solution, the bead was fully submerged in the falling sphere assay and collapsed in the Inversion Assay (Figures 3 and 4).

**Figure 3.**
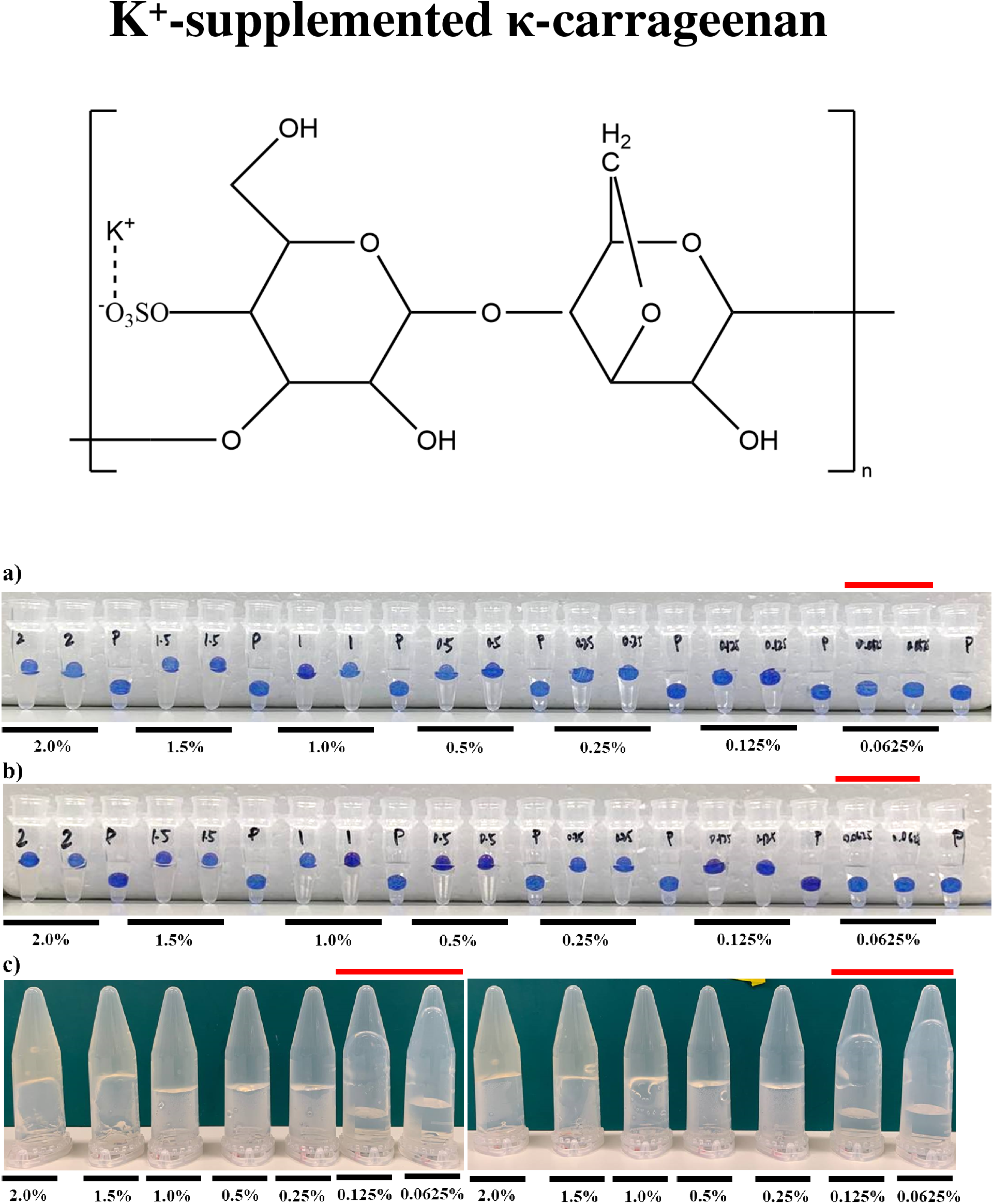
Determining the Minimum Gelling Concentration of potassium-supplemented kappa carrageenan via the Floating Sphere and Inversion Assays. Representation of gelation results of potassium-supplemented kappa carrageenan in (a) Thermocycler – Floating Sphere Assay; (b) Water bath – Floating Sphere Assay and (c) water bath – Inversion Assay. Red line indicates the concentrations below the critical gelling concentration. A minimum of three biological replicates was performed. Results from two additional replicates are presented in Supplementary Figure S3.

**Figure 4.**
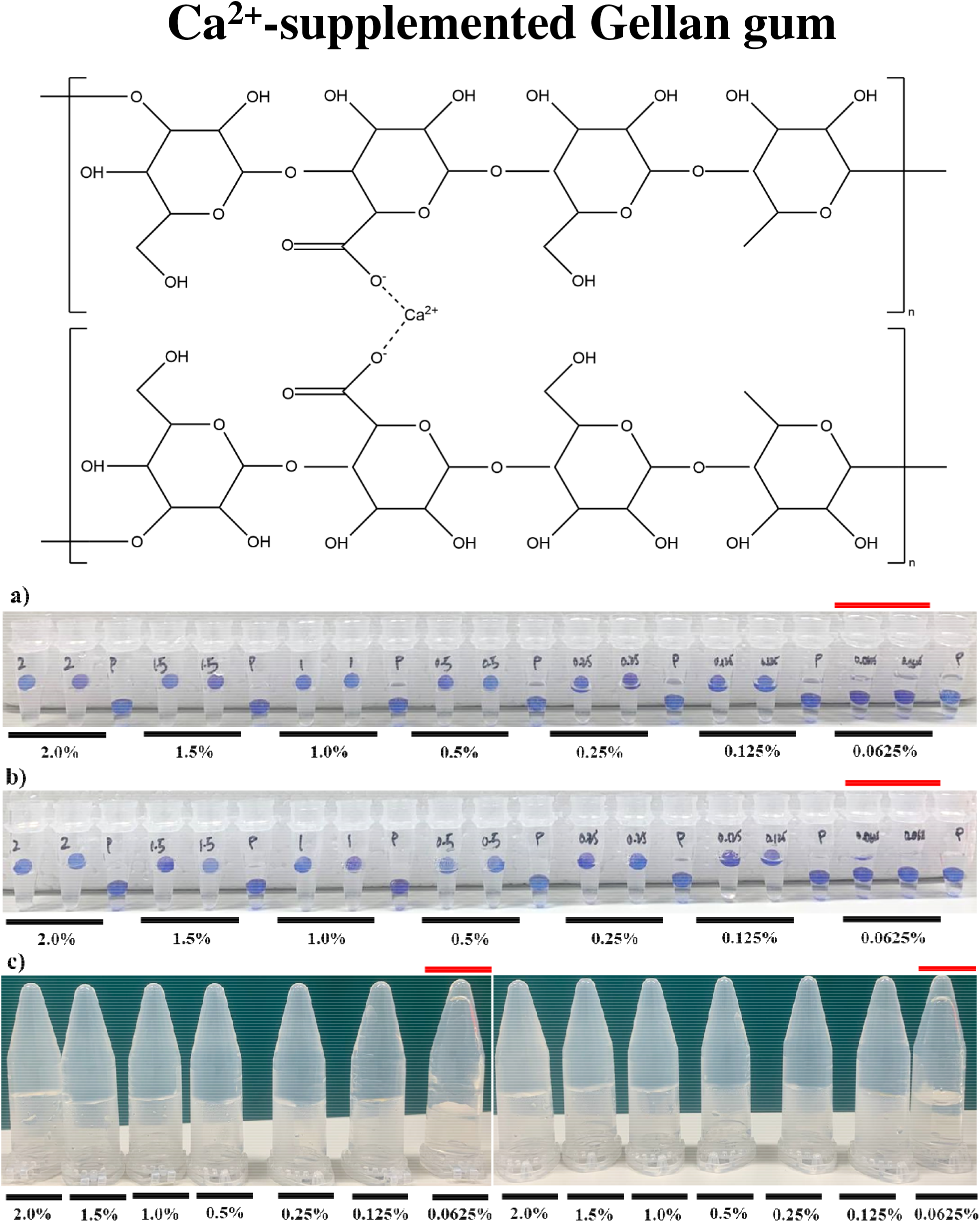
Determining the Minimum Gelling Concentration of calcium-supplemented low acyl gellan gum via the Floating Sphere and Inversion Assays. Representation of gelation results of calcium-supplemented low acyl gellan gum in (a) Thermocycler – Floating Sphere Assay; (b) Water bath – Floating Sphere Assay and (c) Water bath – Inversion Assay. Red line indicates the concentrations below the critical gelling concentration. A minimum of three biological replicates was performed. Results from two additional replicates are presented in Supplementary Figure S4.

Moreover, the values of critical gelling concentrations obtained are largely consistent with values reported in the literature. Critical gelling concentrations was reported to be as low as 0.7% for potassium-supplemented κ-carrageenan solutions (Chen, et al., 2002); 0.1% (Lochhead, 2017) and 0.6% for calcium-supplemented low acyl gellan solutions (J. Tang, Lelievre, Tung, & Zeng, 1994); 0.5% (Pandey, et al., 2019) and 1.0% for gelatin (Djabourov, et al., 1993).

Apart from reduced sample volumes, an additional advantage of the Floating Sphere Assay is its ability to detect the transition zone between strong gels in which the bead floats on top and weak gels in which the bead is partially submerged. This is unlike the conventional Inversion Assay which can only detect the transition zone between gels and non-gelling solutions. This is as reflected from our observations that samples producing weak gels in the Floating Sphere Assay form ‘stable gels’ in the Inversion Assay.

Subsequently, we evaluated the ability of the Floating Sphere Assay to distinguish gels with varying strengths. Several studies have reported that at the same gum concentration in non-calcium supplemented gels, gellan at pH 3.5 would form stronger gels than at pH 7.0. One study reported that rupture stress of the gels at pH 3.5 exceeds that in pH 5.3 and 7.0 counterparts by at least a factor of two (Picone & Cunha, 2011). A separate study observed that at pH 3.5, gellan gum set at much lower concentrations than that of pH 5.2 and, 0.5% gels at pH 3.5 produced a gel as strong as a 2.0% counterpart at pH 5.2 (Cassanelli, Prosapio, Norton, & Mills, 2018). Consistent with these observations, in our Falling Sphere Assay, 1.0% gel corresponds to the transition zone in which beads would be partially submerged in pH 7 gels yet floating at the top in pH 3.5 counterparts (Table 5, Figure 5). Data from two additional biological replicates are presented in Supplementary Figures S5 and S6. Therefore, this indicates that our assay has the sensitivity to distinguish gels with varying strengths. This serves as a cost-effective approach to differentiate gel strength as compared to conventional use of mechanical texture analysers and rheometers (Cassanelli, et al., 2018; Picone, et al., 2011).

**Table 5.**
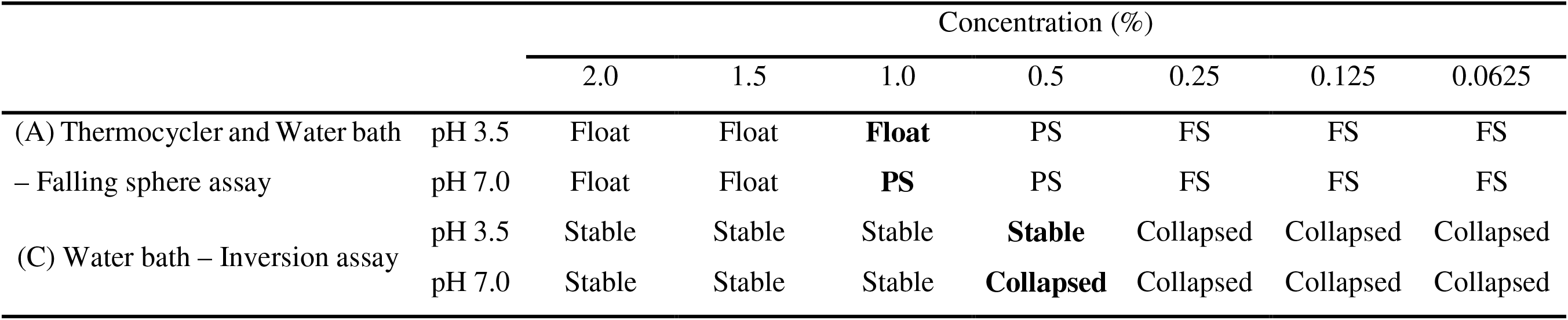
Gelation results of non-supplemented low acyl gellan at pH 3.5 and 7.0.

**Figure 5.**
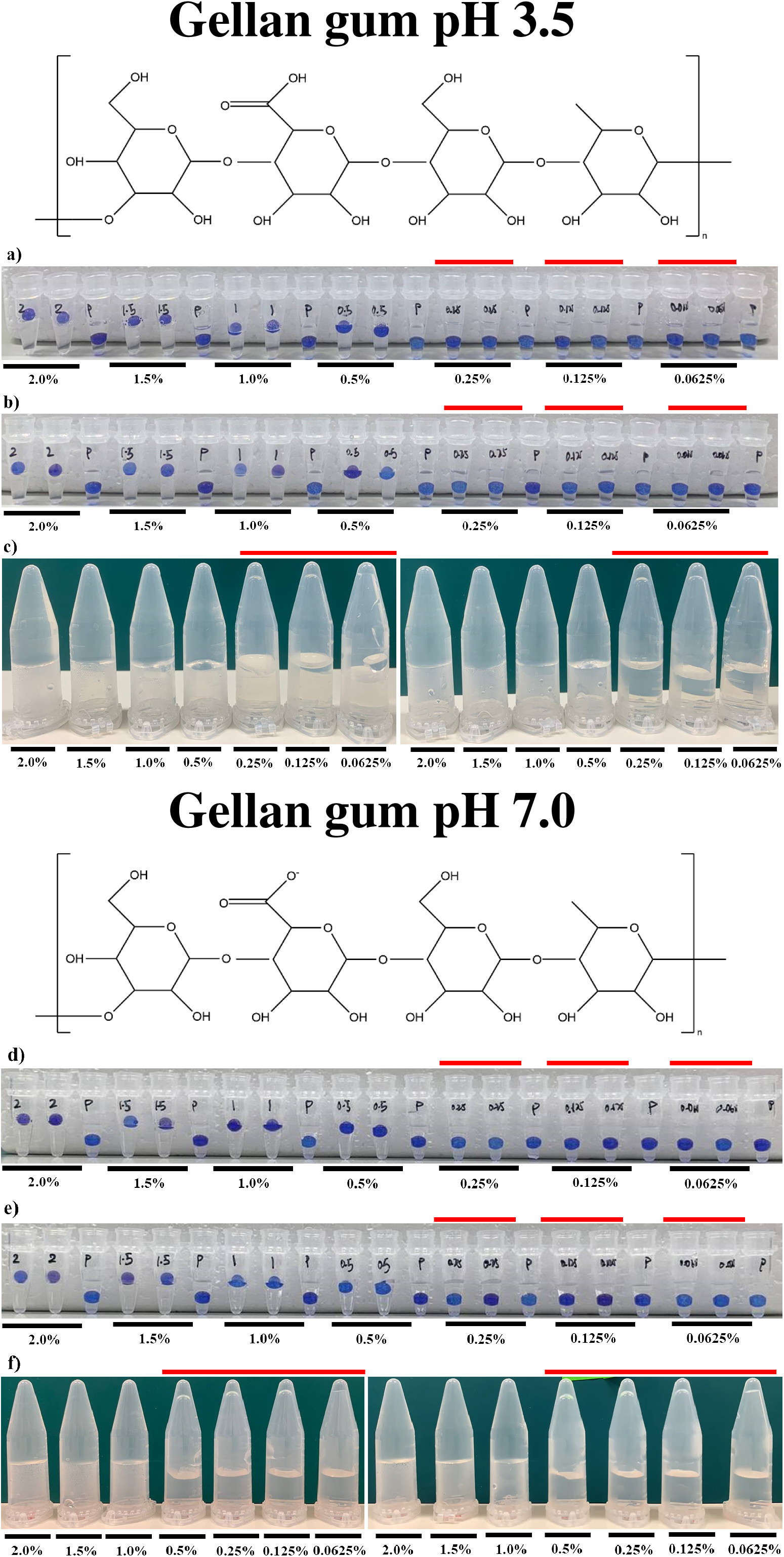
Low acyl gellan gum gels form stronger gels at pH 3.5 compared to pH 7.0. Representation of gelation results of pH 3.5 and pH 7.0 non-supplemented low acyl gellan gum are represented in panels (a)-(c) and (d)-(f) respectively. Panels (a) and (d) Thermocycler – Floating Sphere Assay; Panels (b) and (e) Water bath – Floating Sphere Assay; Panels (c) and (f) Water bath – Inversion Assay. Red line indicates the concentrations below the critical gelling concentration. A minimum of 3 biological replicates was performed. Results from two additional replicates are presented in Supplementary Figure S5.

### Physical principles underlying the Floating Sphere Assay

When the glass bead (density = 2.6 g/cm^3^) is dropped into the solution (density = 1 g/cm^3^) in the PCR tube, it behaves in one of the following three ways. 1. **Bead floats**. In this case, the weight of the bead is supported by the strength of the solid gel generated by the intermolecular and intramolecular interactions involving the solvent, gelling agent and additional buffer components (Figure 6a). 2. **Bead is partially submerged**. In this case, the strength of the gel is unable to completely support the weight of the bead. Although the bead exerted a force exceeding rupture stress (if any formed), strength of the gel in combination with the buoyant force due to submerged portion of the bead, matches the weight on the bead, to keep it transiently afloat (Figure 6b). 3. **Bead is fully submerged**. Weight of the bead exceeds the gel strength (if any formed) and enters the solution (Figure 6c).

**Figure 6.**
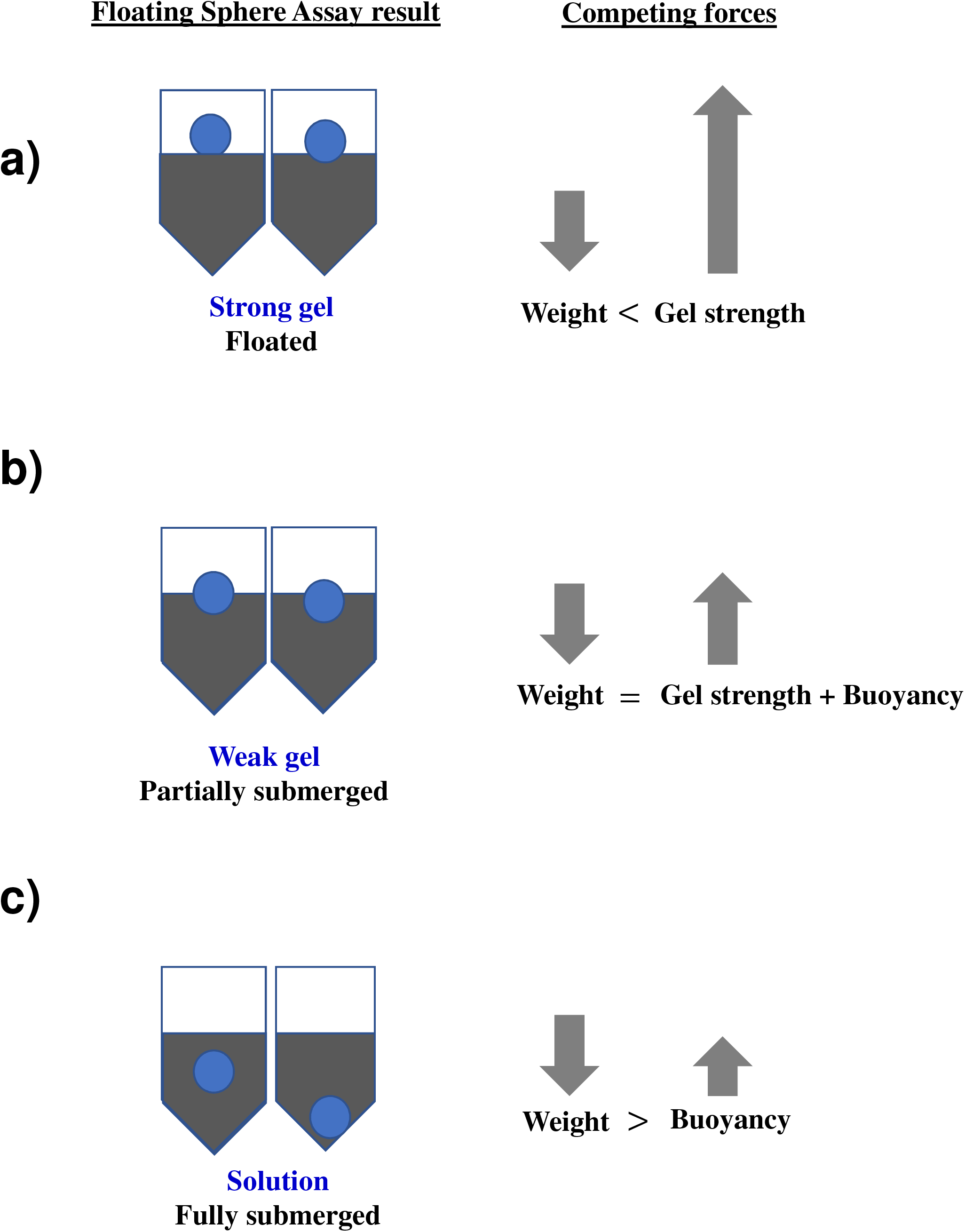
Physical principles underlying the Floating Sphere Assay. Forces acting on the bead for the three possible outcomes of the Floating Sphere assay. (a) Bead is floating; (b) bead is partially submerged; (c) bead is completely submerged. Please see the text for details.

## 7 Limitations and future work

Our falling sphere assay is technically simple, economical and can be easily adapted for high-throughput screening for gel formation. It might be difficult to distinguish very viscous solutions from weak gels. To differentiate between weak gels and viscous solutions when the bead is partially submerged, lower density beads could be used. This is because the weak gels are more likely to support beads of lower density. Correspondingly, to differentiate strength amongst gels when bead is floating at the top, higher density beads (e.g., steel) might be useful.

Moreover, this assay could be potentially adapted to facilitate development of a miniaturized texture profile analyser. This would require improved precision of the Floating Sphere Assay, achievable through automating the position at which the ball is deposited into the sample tube. This would serve to standardize the height and angle from which the sphere is dropped, thereby controlling the velocity and area by which the glass bead contacts the surface of the gel solution. This would serve to standardize the depth of depression/compression should the sample be a solid gel and, depth of penetration should the sample be in a liquid-state. Microscopic imaging of the former serves can serve as a means to characterize springiness of strong gels (i.e. recovery on compression by a fixed force) while the latter can differentiate viscosity of weaker gels. Lastly, dimensions of the ball used could be further reduced to facilitate the use of even smaller sample volumes to detect gel formation.

## 8 Conclusion

The Floating Sphere Assay is a reliable miniaturised method for gelation with high accuracy to detect MGC and sensitivity to differentiate between strong and weak gels. This can be utilized in high-throughput screens for novel protein gelling agents derived through recombinant or natural extraction approaches.

## Acknowledgements

This research was supported by the Singapore Food Story Seed grant awarded to PA. UJT is funded by the A*STAR Graduate Scholarship. JCWH is supported by the A*STAR Research Internship programme.

## Author contributions

**Uma Jingxin Tay:** Investigation Formal analysis and Writing-Original draft

**Megan Goh:** Investigation Formal analysis and Writing-Review and editing

**Jeralyn Ching Wen Hui:** Investigation Formal analysis and Writing-Review and editing

**Prakash Arumugam:** Conceptualization, Supervision and Writing-Review and editing

## Supplementary materials

**Legends for Supplementary Figures**

**Supplementary Figure S1.**
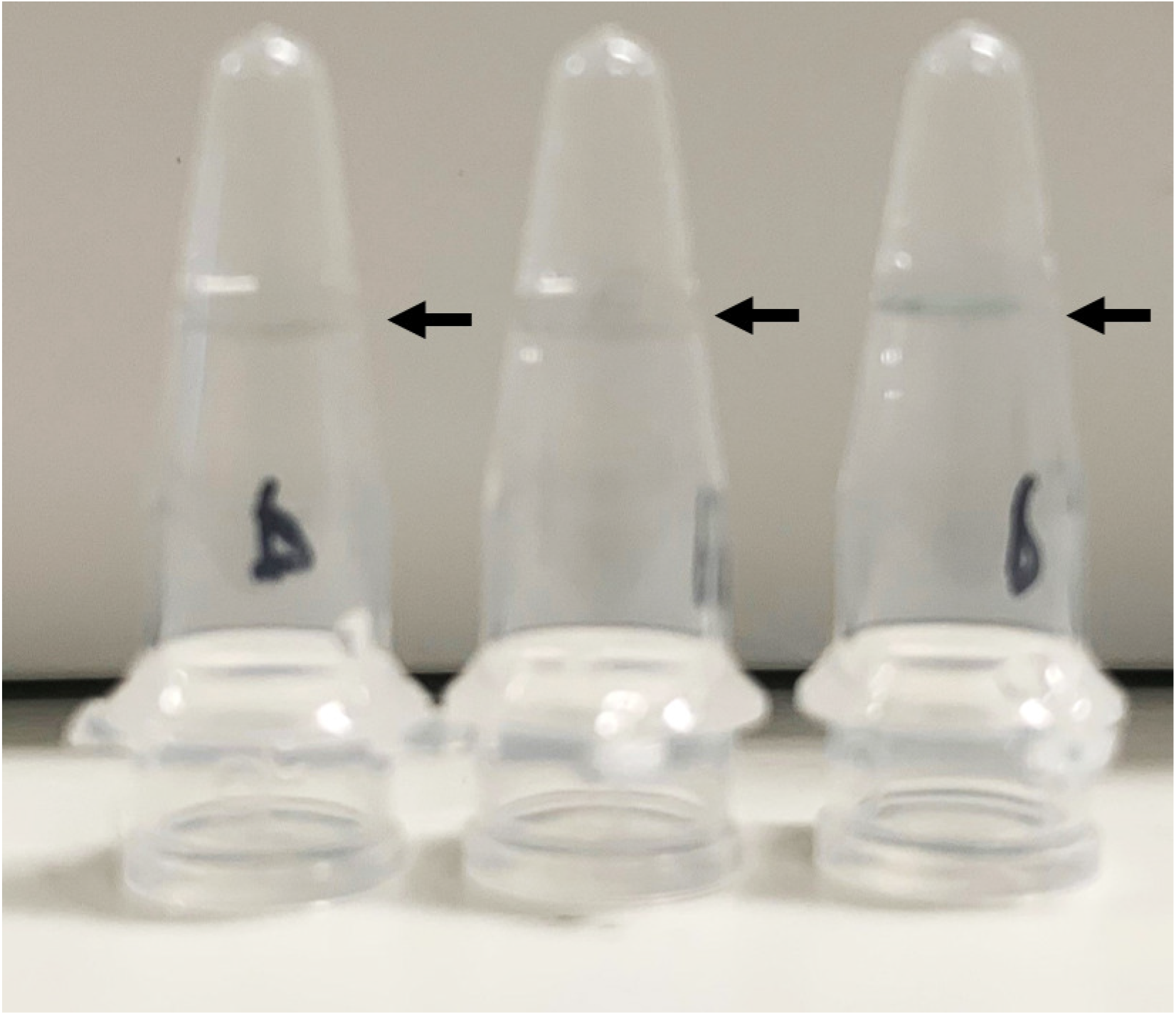
Inverted PCR tubes containing PBS solution. Solution level is marked by black arrows.

**Supplementary Figure S2.**
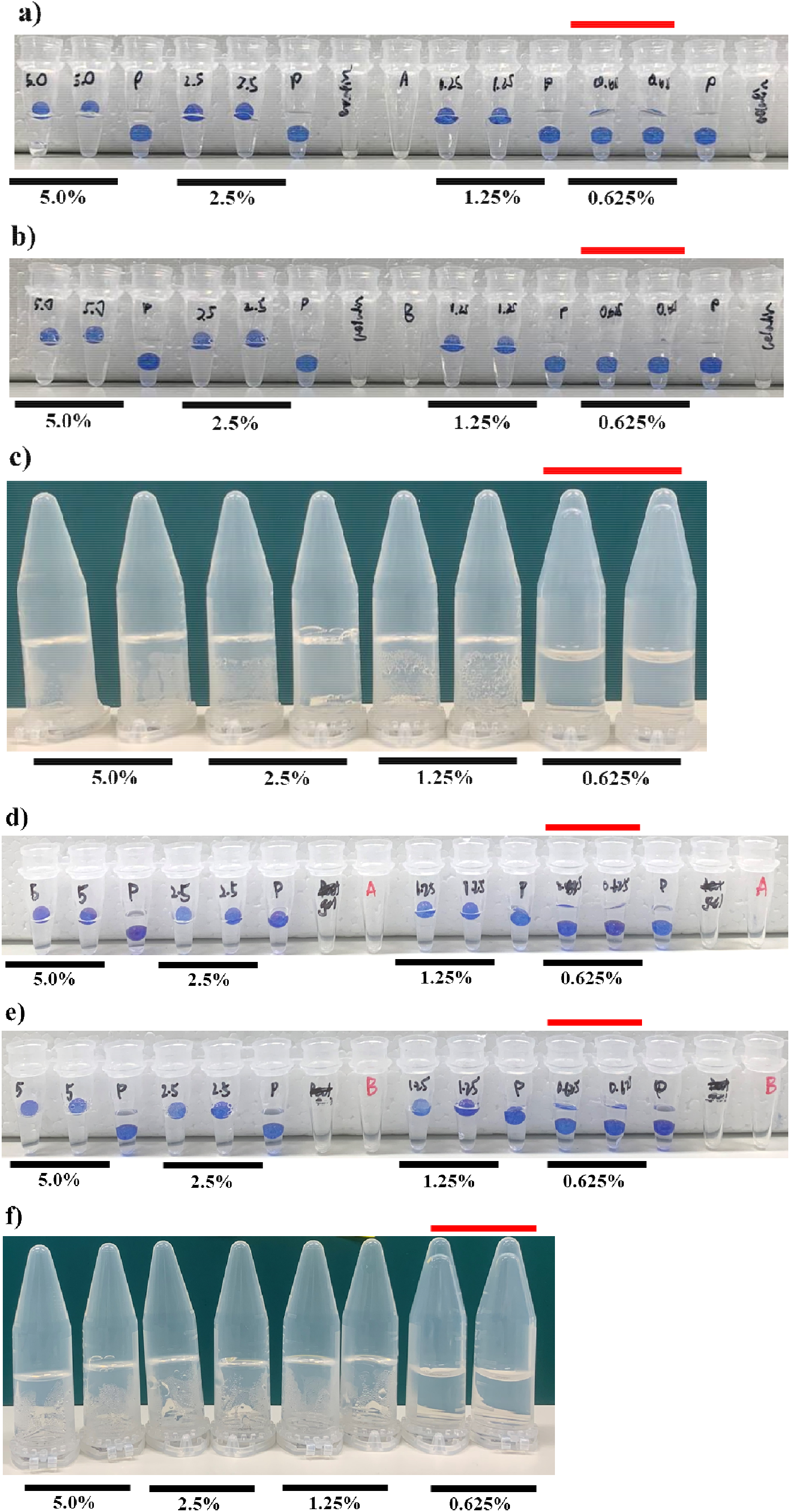
Data from two additional replicates for gelation results of gelatin are presented in (a) – (c) and (d) – (f) respectively. (a) and (d) correspond to Thermocycler – Floating Sphere Assay; (b) and (e) correspond to Water bath – Floating Sphere Assay; (c) and (f) correspond to Water bath – Inversion Assay.

**Supplementary Figure S3.**
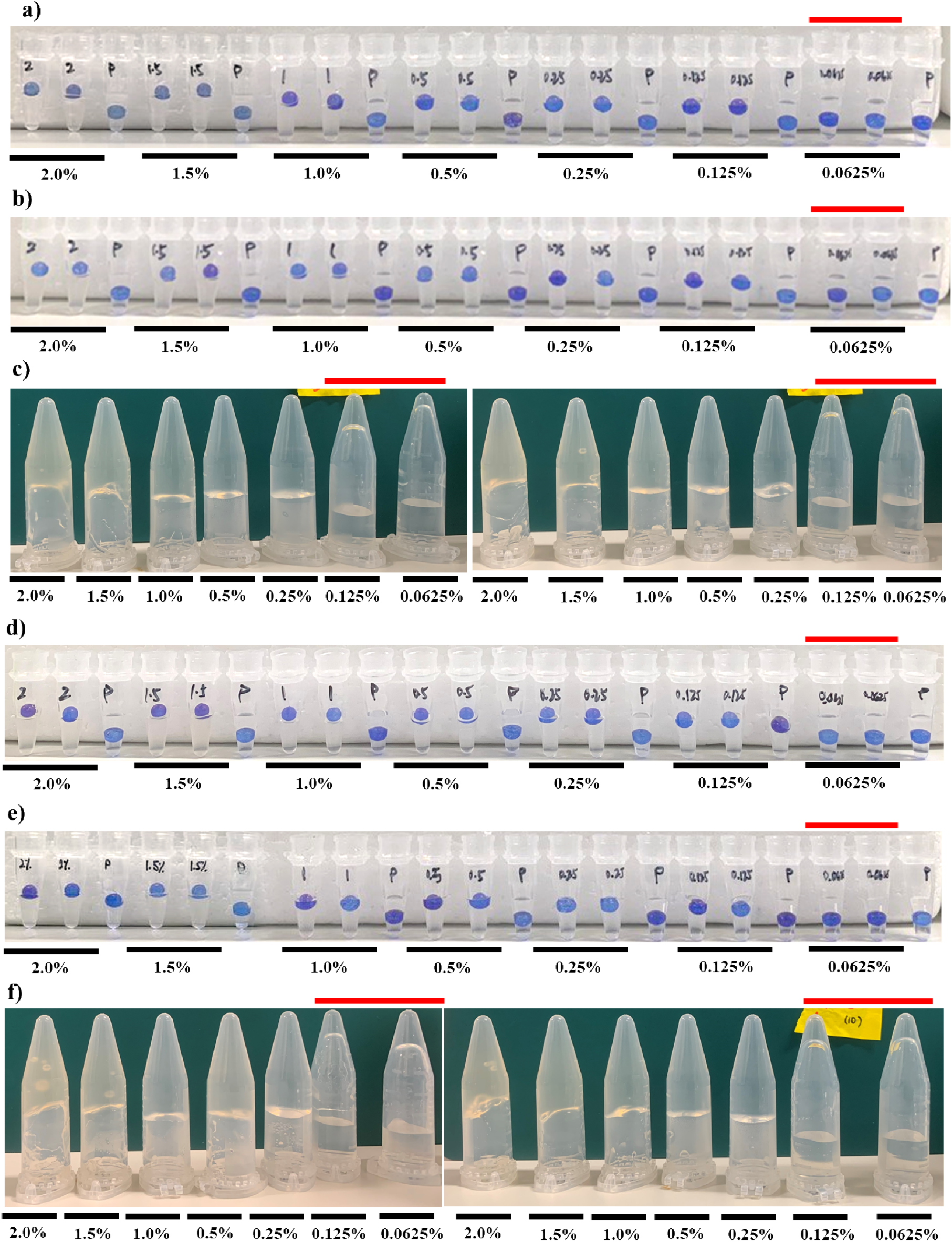
Data from two additional replicates for gelation results of potassium-supplemented kappa carrageenan are presented in (a) – (c) and (d) – (f) respectively. (a) and (d) correspond to Thermocycler – Floating Sphere Assay; (b) and (e) correspond to Water bath – Floating Sphere Assay; (c) and (f) correspond to Water bath – Inversion Assay.

**Supplementary Figure S4.**
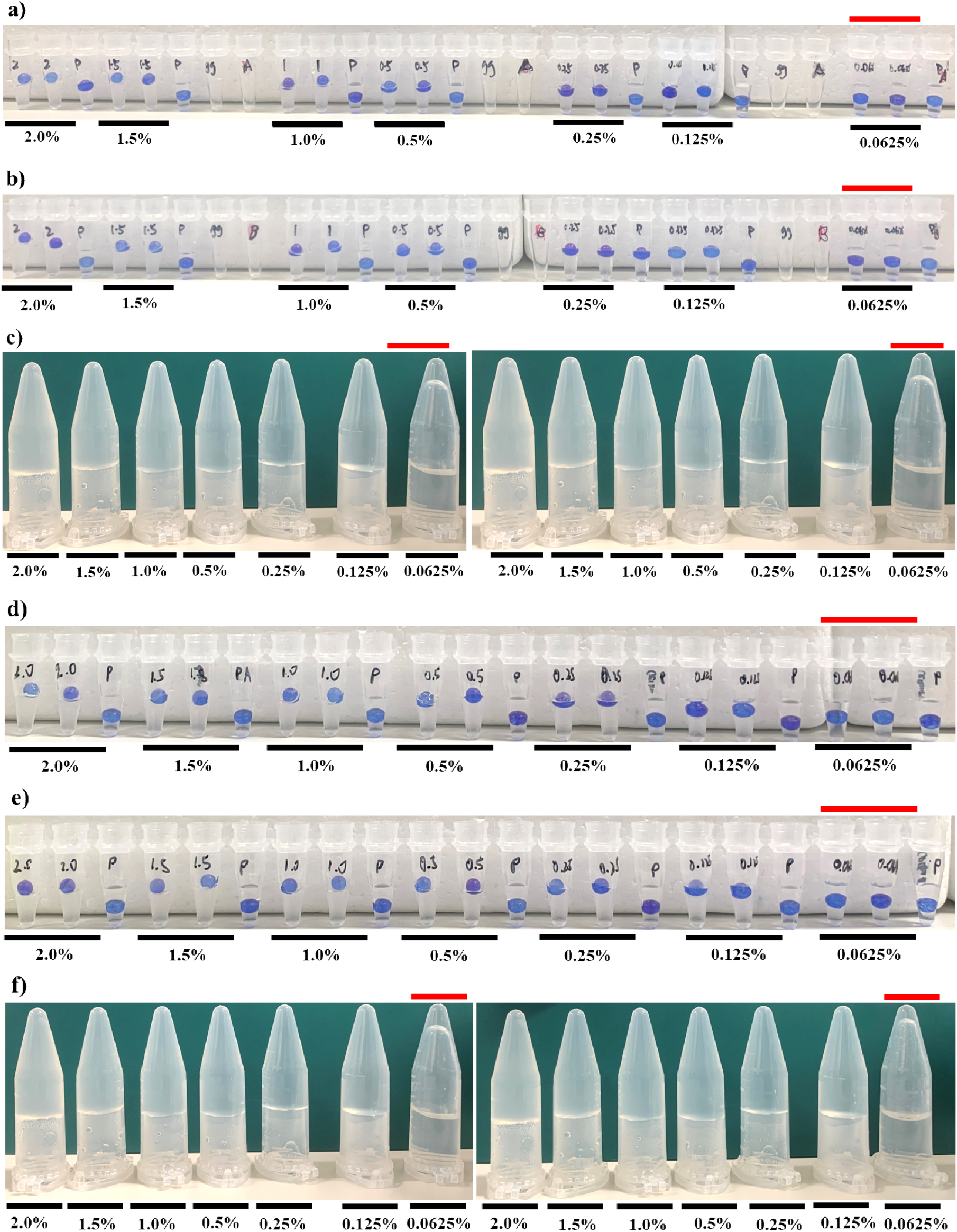
Data from two additional replicates for gelation results of calcium-supplemented low acyl gellan gum are presented in (a) – (c) and (d) – (f) respectively. (a) and (d) correspond to Thermocycler – Floating Sphere Assay; (b) and (e) correspond to Water bath – Floating Sphere Assay; (c) and (f) correspond to Water bath – Inversion Assay..

**Supplementary Figure S5.**
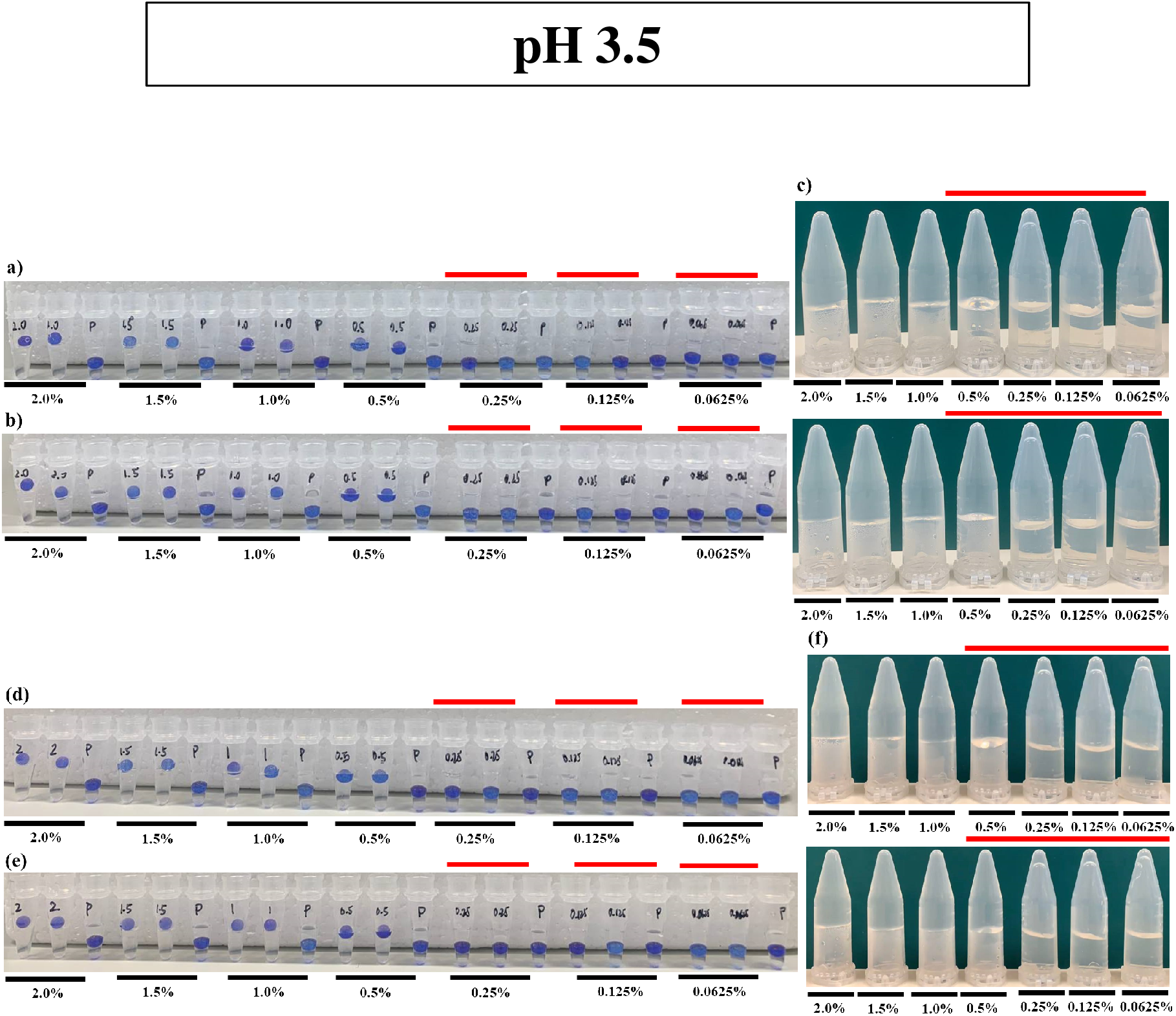
Data from two additional replicates for gelation results of non-supplemented low acyl gellan gum at pH 3.5 are presented in (a) – (c) and (d) – (f) respectively. (a) and (d) correspond to Thermocycler – Floating Sphere Assay; (b) and (e) correspond to Water bath – Floating Sphere Assay; (c) and (f) correspond to Water bath – Inversion Assay.

**Supplementary Figure S6.**
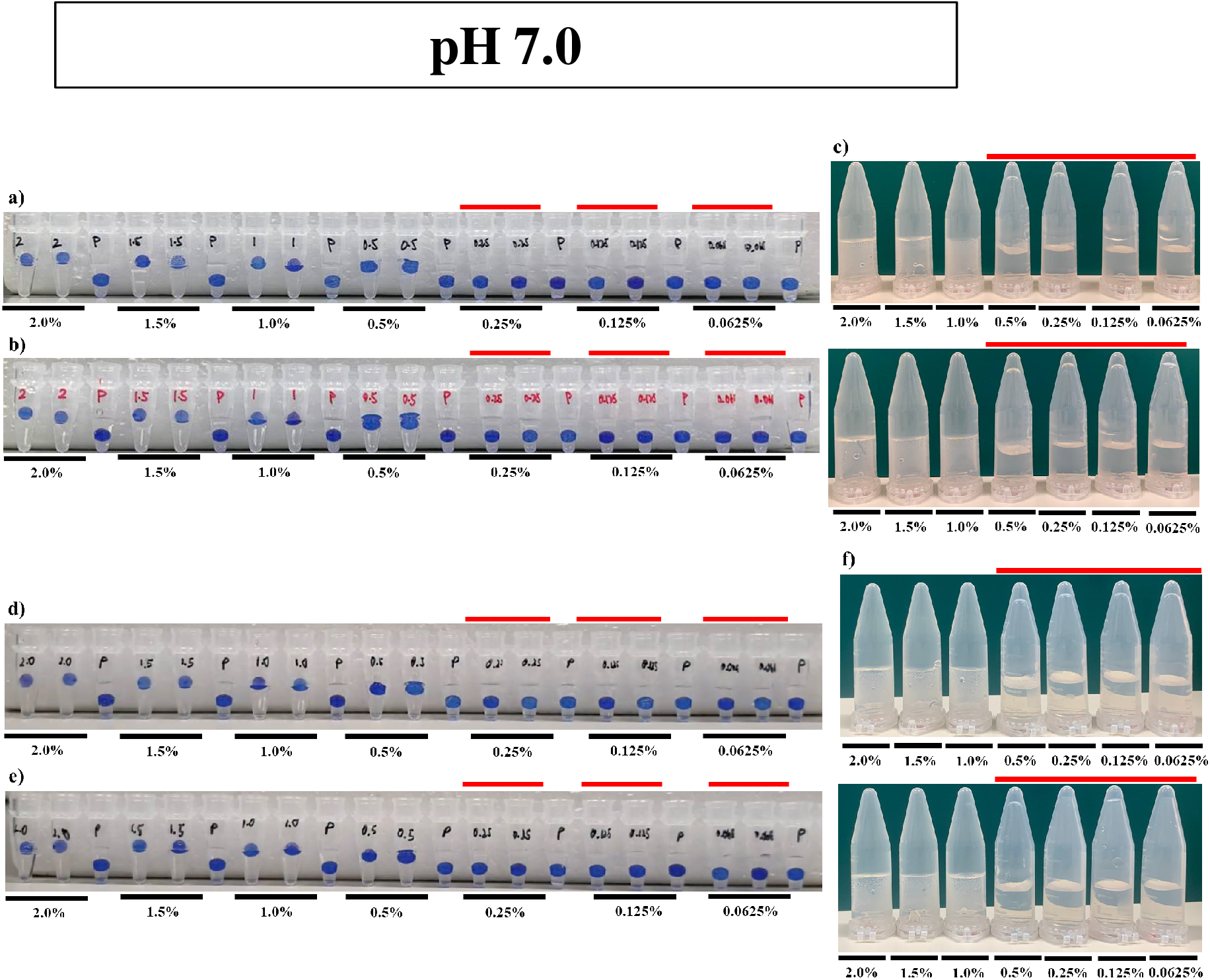
Data from two additional replicates for gelation results of non-supplemented low acyl gellan gum at pH 7.0 are presented in (a) – (c) and (d) – (f) respectively. (a) and (d) correspond to Thermocycler – Floating Sphere Assay; (b) and (e) correspond to Water bath – Floating Sphere Assay; (c) and (f) correspond to Water bath – Inversion Assay.

